# Increased excitation enhances the sound-induced flash illusion by impairing multisensory causal inference in the schizophrenia spectrum

**DOI:** 10.1101/2024.05.29.596551

**Authors:** Renato Paredes, Francesca Ferri, Vincenzo Romei, Peggy Seriès

## Abstract

The spectrum of schizophrenia is characterised by an altered sense of self with known impairments in tactile sensitivity, proprioception, body-self boundaries, and self-recognition. These are thought to be produced by failures in multisensory integration mechanisms, commonly observed as enlarged temporal binding windows during audiovisual illusion tasks. To our knowledge, there is an absence of computational explanations for multisensory integration deficits in patients with schizophrenia and individuals with high schizotypy, particularly at the neurobiological level. We implemented a multisensory causal inference network to reproduce the responses of individuals who scored low in schizotypy in a simulated double flash illusion task. Next, we explored the effects of recurrent excitation, cross-modal and feedback weights, and synaptic density on the visual illusory responses of the network. Using quantitative fitting to empirical data, we found that an increase in the weights of the recurrent excitatory connectivity in the network enlarges the temporal binding window and increases the overall proneness to experience the illusion, matching the responses of individuals scoring high in schizotypy. Moreover, we found that an increase in excitation increases the probability of inferring a common cause from the stimuli. We propose an E/I imbalance account of reduced temporal discrimination in the SCZ spectrum and discuss possible links with Bayesian theories of schizophrenia. We highlight the importance of adopting a multisensory causal inference perspective to address body-related symptomatology of schizophrenia.

## 1 Introduction

Individuals with schizophrenia show a disturbed sense of self, with impairments in somatosensory processing such as tactile sensitivity (Chang and Lenzenweger, 2001, 2004, 2005) and proprioception (Michael and Park, 2016; Thakkar et al., 2011). They also have deficits in higher-order processes essential for a coherent sense of self, like body ownership (Rossetti et al., 2020; Costantini et al., 2020; Zopf et al., 2021; He et al., 2022), bodily-self boundary (Park et al., 2009; Holt et al., 2015; Di Cosmo et al., 2018, 2021), and self-recognition (Ferroni et al., 2019; Sandsten et al., 2020). These differences often stem from disruptions in multisensory integration, the neural process that combines sensory information to form a unified perceptual experience (Stein and Stanford, 2008; Stein et al., 2010). Effective interaction with the world depends on integrating vestibular, proprioceptive, and interoceptive information with external sensory data, making multisensory integration crucial for self-coherence and consciousness (Noel et al., 2018b; Serino, 2019).

Impairments in multisensory integration in the schizophrenia spectrum are also observed with non-body-related stimuli, as audiovisual illusion experiments suggest. Patients with schizophrenia are more prone to experience the streaming-bouncing illusion (Zvyagintsev et al., 2017), the McGurk effect (Martin et al., 2013) and the double flash illusion (Haß et al., 2017). These differences have also been observed in high schizotypal individuals for the McGurk (Muller et al., 2021) and the double flash illusion (Ferri et al., 2018). Despite the amount of empirical evidence available, there is as yet no theoretical understanding of why multisensory integration is altered in the schizophrenia spectrum.

These generalised failures in multisensory integration mechanisms could be explained by the enlarged temporal binding window (i.e., the epoch of time in which stimuli from different modalities are more likely to be perceptually bound) (TBW) observed in the schizophrenia spectrum in several paradigms (Ferri et al., 2017; Zhou et al., 2018; Dalal et al., 2021; Di Cosmo et al., 2021). In general, experiments that evaluate the judgment of the relative timing of stimuli with different onset asynchronies reveal that patients with schizophrenia and individuals with high schizotypy show a longer TBW compared to controls (Zhou et al., 2018; Dalal et al., 2021; Di Cosmo et al., 2021; Fotia et al., 2021).

In the double flash illusion (DFI) paradigm, schizophrenia patients more frequently perceived an illusory flash and had a broader temporal window of integration (TWI) for auditory and visual stimuli. This is likely due to the increased cross-modal influence of the second auditory input onto the representation of the visual input, which is then perceived to be repeated, even at longer sound onset asynchronies (Haß et al., 2017; Cooke et al., 2019). Similarly, high-schizotypy individuals showed an enlarged TWI and greater susceptibility to the illusion than those with low schizotypy, with both measures positively correlated (Ferri et al., 2018).

In general, audiovisual illusions have been consistently accounted for in healthy individuals with causal inference models of multisensory integration (Cao et al., 2019; Kayser and Shams, 2015; Magnotti et al., 2018; Mohl et al., 2020; Rohe et al., 2019). These models mainly use the Bayesian inference framework (Körding et al., 2007), which provides a high-level description (i.e. computational level of analysis according to Marr (2010)) of the computations carried out by the brain to integrate unisensory signals. To our knowledge, biologically plausible neural network models of multisensory causal inference (Cuppini et al., 2017; Fang et al., 2019; Rideaux et al., 2021) have not yet been used to explore the neural mechanisms behind the proneness to experience audiovisual illusions in individuals in the schizophrenia spectrum.

In particular, a comprehensive explanation of why patients with schizophrenia and individuals with schizotypal traits are more prone to experience audiovisual illusions is lacking. Similarly, there is no clear mechanistic understanding of how an enlarged TBW (Ferri et al., 2017; Zhou et al., 2018; Dalal et al., 2021; Di Cosmo et al., 2021) is related to failures in multisensory integration/segregation and causal inference in the schizophrenia spectrum (for a current debate, see Samaha and Romei (2024) and Schoffelen et al. (2024)). Moreover, it is unclear how impairments in these neural processes are related to bodily self-aberrations (Costantini et al., 2020; Zopf et al., 2021; Holt et al., 2015; Di Cosmo et al., 2018; Sandsten et al., 2020) and symptoms of schizophrenia and schizotypal traits.

To this end, we build on recent advances in multisensory causal inference network models in healthy individuals (Cuppini et al., 2017) to reproduce the greater proneness to double flash illusion observed in individuals with high schizotypy. Our model is based on the idea that inferring a common cause to audiovisual stimuli increases the occurrence of the DFI (Hirst et al., 2020). In particular, we evaluate the hypothesis that an imbalance of E/I in sensory networks increases temporal binding windows (Ferri et al., 2017), which in turn alters causal inference.

## 2. Methods

### 2.1. Experimental task

Our computational model simulates the double flash illusion (DFI) experiment carried out in individuals with high schizotypy (Ferri et al., 2018) (see Figure 1 for an illustration). The experiment consisted of presenting one visual flash (lasting for 12 ms) to the participants accompanied by two tones (lasting for 7 ms each). The first tone was always displayed simultaneously with the flash, whereas the second tone could be presented at 15 different stimulus onset asynchronies (SOA) from the first tone (ranging from 36 to 204 ms). The experiment consisted of 150 trials, and the second tone was presented 10 times for each SOA.

**Figure 1:**
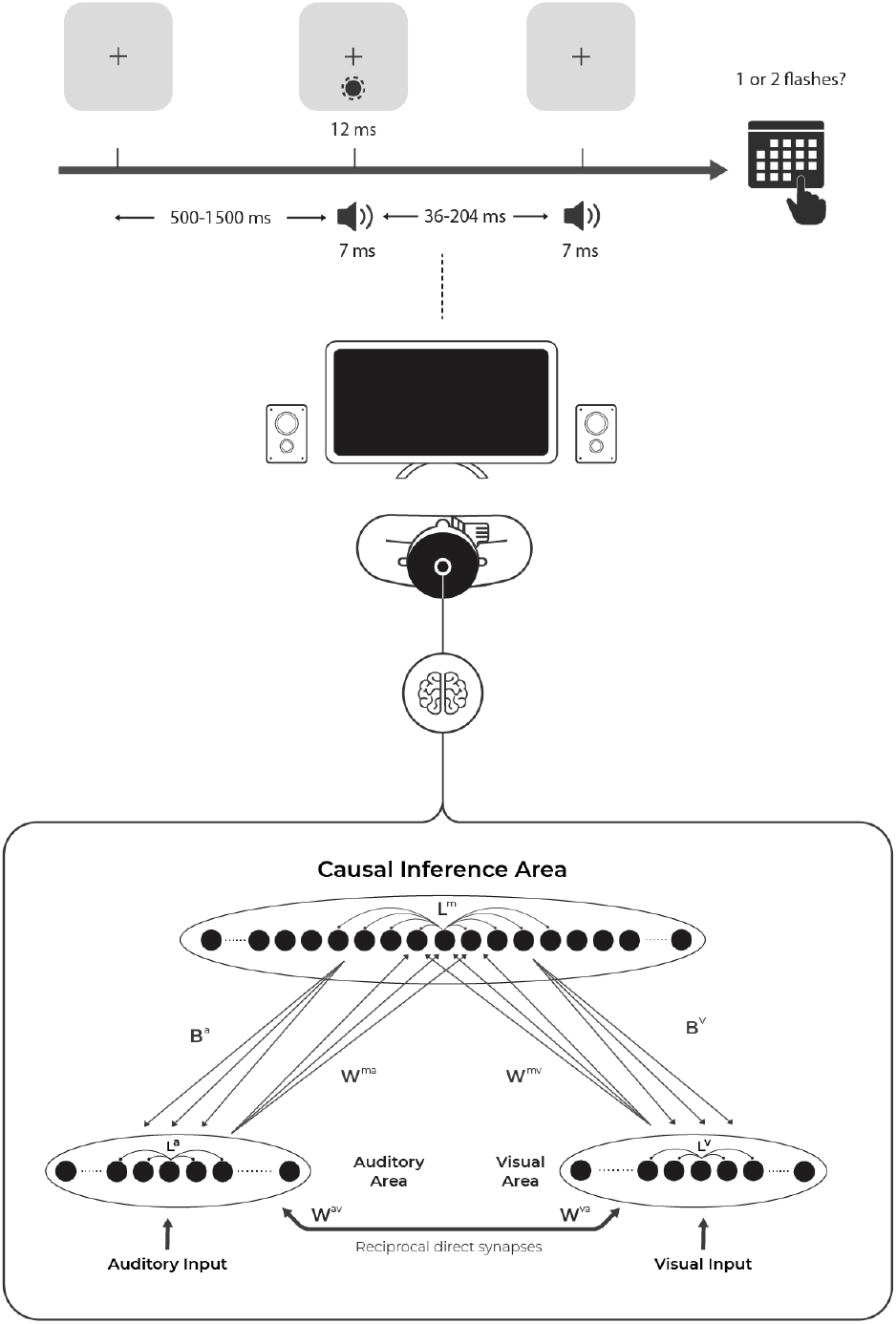
Double flash illusion experimental paradigm and Multisensory Causal Inference network model. In this experimental setup, two speakers and a monitor are placed in front of the participant. The black arrow indicates the evolution of the presentation of stimuli in time: one visual flash (lasting 12 ms), two tones (lasting 7 ms each), and an interval (randomly ranging from 500 to 1500 ms). The first tone was always displayed simultaneously with the flash, whereas the second tone could be presented at 15 different stimulus onset asynchronies (SOA) from the first tone (ranging from 36 to 204 ms). In each trial, participants were asked to verbally report whether they perceived one or two visual flashes. The shorter the SOA, the more likely a second illusory visual flash is perceived. The network model is composed of two unisensory areas (auditory and visual) connected with a multisensory causal inference area. These areas are arranged to encode a portion of 30^°^ of the external space.

In each trial, participants were asked to verbally report whether they perceived one or two visual flashes. They are asked to focus only on visual stimuli and ignore tones (see Ferri et al. (2018) for an extensive description). The manipulation of the delay of the second auditory stimulus produces the subjective experience that the shorter the delay, the more likely a second illusory visual flash is perceived (Cecere et al., 2015; Shams et al., 2002; Cooke et al., 2019). In the original study, the overall proneness to the illusion (OPI) and the temporal window of illusion (TWI) were compared in two groups separated by their scores on the Schizotypal Personality Questionnaire (SPQ) (Ferri et al., 2018).

### 2.2. Multisensory Causal Inference network model of DFI

We model the DFI by extending the audiovisual network model proposed by Cuppini et al. (2014). Following Cuppini et al. (2017), the main modification is the incorporation of a multisensory causal inference layer with feedforward and feedback connectivity to unisensory layers. This multisensory layer implements a mechanism for causal inference by determining the likelihood that the presented sensory stimuli (flash and beeps) originate from a common source versus independent sources. Importantly, this structural change allows us to model the DFI as part of a causal inference process described at the neural level.

In short, the model describes two reciprocally connected unisensory areas (auditory and visual) connected with a multisensory area that computes causal inference (see Figure 1 for an illustration). Each of the three areas consists of 30 rate neurons that encode 30° of external space (see Figure 1 for an illustration). The neurons within each area are connected through lateral synapses following a Mexican hat pattern of excitation and inhibition (*L*^*a*^, *L*^*v*^, *L*^*m*^). Our network uses a “minimal” spatial representation, extending beyond the stimulus to incorporate recurrent connectivity and to be able to capture spatiotemporal relationships.

Our model includes cross-modal connectivity, as supported by modelling studies on audio-visual integration (Cuppini et al., 2014, 2017; Ursino et al., 2019) and empirical evidence obtained from research on primates (Eckert et al., 2008) and humans (Werner and Noppeney, 2010; Gurtubay-Antolin et al., 2021). Neurons in the auditory and visual areas are connected to neurons of the other unisensory area through cross-modal synapses *W*^*av*^, *W*^*va*^.

The neurons in the multisensory area are connected to the neurons in the auditory and visual areas that encode the proximal portions of the space, both through feedforward (*W*^*mv*^, *W*^*ma*^) and feedback synapses (*B*^*vm*^, *B*^*am*^). All the inter-areal synapses are symmetrical and followed a Gaussian pattern (see the Supplementary Information for an extensive mathematical description of the model).

Following Rohe and Noppeney (2015b), our network model incorporates two consecutive hierarchical levels of multisensory processing. In the unisensory areas, the model computes the spatiotemporal location of external stimuli (solving the so-called “implicit”^1^ causal inference problem). In the multisensory area, the model evaluates whether stimuli originate from a single source (solving the “explicit” causal inference problem), mimicking the responses of neurons in parietal-temporal association cortices. The network also features feedback connectivity, enabling iterative refinement of causal inferences across the network (Rohe et al., 2019; Cao et al., 2019).

Our model does not account for any noise in the representation of the inputs or neural activities. We did not conduct an exhaustive comparison of all conceivable network architectures that could potentially explain the DFI, as such an analysis lies beyond the scope of this study. To test whether the causal inference area was necessary in the model, we simulated an architecture without it and found that it could not account for response performances (see dashed lines in Figure 4A). An extensive description of the model and values of the parameters can be found in the Supplementary Information.

### 2.3. Experiment simulation in the network

In simulating audiovisual stimuli in the DFI, the network receives Gaussian 1-D inputs centred in the encoded space. Visual inputs last 12 ms, auditory 7 ms. Visual and first auditory stimuli start 16 ms after trial onset. The second auditory stimulus starts at 15 delays ranging from 36 to 204 ms after the first stimulus, with the network running for 600 ms for each delay. Given the deterministic nature of the model, each experimental condition is only simulated once.

We model DFI as a multisensory integration process with early and late cross-modal interactions in distinct cortical hierarchies (Hirst et al., 2020; Keil, 2020). The literature informs that an early interaction arises from cross-modal input between primary sensory cortices observed between 35-90 ms after the onset of the stimuli. These have been observed in Heschl’s gyrus and Calcarine cortex (Shams et al., 2005; Raij et al., 2010), and functionally supported by TMS and EEG evidence (Romei et al., 2007; Cappe et al., 2010; Romei et al., 2012). Next, a delayed interaction arises from the input of feedforward-feedback connectivity with higher cortical regions (e.g. angular gyrus and the superior temporal sulcus) observed in the 130-600 ms post-stimulus range (Mishra et al., 2007; Balz et al., 2016a; Keil and Senkowski, 2018; Keil, 2020). Critically, brain activity consistent with Bayesian Causal Inference^2^ has been observed in anterior parietal areas in the 200-400 ms range (Rohe et al., 2019).

To approximate these temporal dynamics in the network model, we consider a temporal delay of 16 ms for cross-modal interactions (as in Cuppini et al. (2014)) and 95 ms for feedforward and feedback interactions. We established these temporal delays to model the empirical observations that crossmodal delays (typically around 35-90 ms) are considerably lower than feedforward-feedback delays (typically around 130-600 ms) (Keil and Senkowski, 2018; Keil, 2020). Moreover, following Cuppini et al. (2014), we included temporal filters for auditory, visual, and multisensory neurons to mimic the temporal evolution of neural input and synaptic dynamics. Such filters consider specific time constants that define the temporal dynamics of each group of neurons (auditory, visual, or multisensory).

The responses of the participants in the DFI experimental paradigm (Ferri et al., 2018) can be simulated by a principled readout of the unisensory activity of the network. Visual neurons’ peak activity values above 0.15 (where activity is normalised from 0 to 1) were read out as the probability of perceiving a flash. Consistent with the methodology delineated in Cuppini et al. (2017), this threshold was established to indicate that a neuron must exhibit activity up to 15% of its maximal firing rate to achieve a distinguishable perception. The probability of perceiving two flashes within a single trial was defined as the outcome of a combined probability operation: the product of the probability of detecting an initial flash and the probability of detecting a subsequent flash, contingent upon the condition that these probability values exceed the established detection threshold and the valley between them is at least .15 in depth^3^.

To reproduce the sigmoid fit of the average responses of the low schizotypy group (Ferri et al., 2018), we fitted the parameters that define the crossmodal, feedforward and feedback synaptic weights, as well as the auditory, visual, and multisensory time constants (see Supplementary Information for a full description of the fitting procedure). The purpose of this manipulation was to calibrate the model to reproduce responses that closely resemble those observed in healthy individuals in the empirical study. Hence, in the following this network setup will be named the low schizotypy (L-SPQ) model and will be taken as the baseline for further exploration.

An illustration of the dynamics of the baseline model is shown in Figure 2. For the network readout, we selected the neuron with the maximal response to the stimuli delivered in each layer (winner take all mechanism). We observe that at every SOA, the visual neuron shows two peaks of activity: the first generated by auditory and visual stimuli, and the second as a result of feedback neural input. This feedback input depends on the degree of activation of the multisensory causal inference neuron, which in turn depends on the combined feedforward input from auditory and visual neurons. The dynamics of the model show that the closer the auditory stimuli in time (lower SOA), the higher the combined feedforward signal, as a consequence of cross-modal neural input.

**Figure 2:**
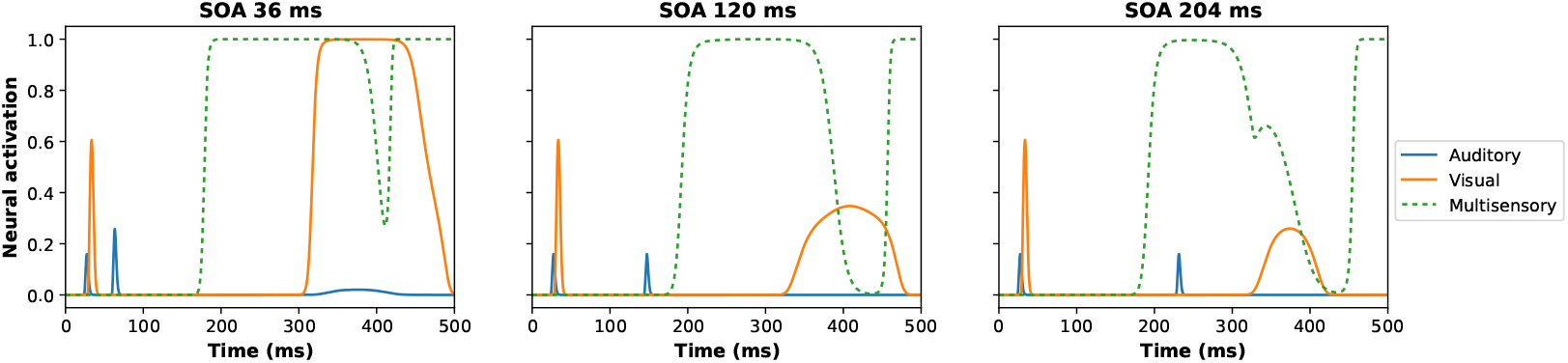
Base model (L-SPQ) dynamics. The network receives a 12 ms visual input and a 7 ms auditory input simultaneously, followed by a second auditory input at 15 different stimulus onset asynchronies (SOAs) from 36 to 204 ms. The figure shows the activity of auditory (blue line), visual (orange line) and multisensory (dashed green line) neurons. The auditory neuron shows two peaks of activity generated by the auditory stimuli received by the network. The visual neuron shows two peaks of activity: the first generated by the visual stimulus received by the network, and the second as a result of neural feedback. The multisensory neuron shows one activity peak at lower SOAs (36 ms) and two distinct peaks at larger SOAs (120 and 204 ms). This pattern results from feedforward input from auditory and visual neurons, where closer auditory stimuli increase the combined signal due to cross-modal neural interactions. Consequently, the greater and more sustained the activity of the multisensory neuron, the stronger the feedback input to the unisensory areas, resulting in an elevated peak of illusory visual activity. By design, feedforward input coming from unisensory activity is delayed by 95 ms before reaching the multisensory area.

Given these model dynamics in which the multisensory neuron tends to reach its maximal firing rate at every SOA, the multisensory neurons’ activity above .8 was read out as the probability of inferring that the visual input and the second auditory input were caused by a single cause (peak activity values below this threshold were assigned a probability of zero). The probability of inferring a single cause from the stimuli within a single trial was calculated as the complement of the probability of inferring two different causes (i.e. the product of two peak values of multisensory neurons, contingent upon the condition that these are above the detection threshold and the valley between them is at least .80 in depth).

### 2.4. Modelling the influence of high schizotypy in the network

We evaluated the potential consequences of four observed impairments in the SCZ spectrum on the sound-induced flash illusion: excitation/inhibition (E/I) imbalance (Jardri et al., 2016; Leptourgos et al., 2022), early crossmodal processing deficits (Magnée et al., 2009; Balz et al., 2016b), failures in top-down signalling (Sterzer et al., 2018), and synaptic density decrease (Ellison-Wright and Bullmore, 2009).

#### 2.4.1. Modulation of the E/I balance

The modulation of the E/I balance was implemented by increasing or decreasing the strength of the recurrent excitatory connectivity. This was done by modifying the value of the parameter 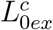 in each area, defined in equation 1:

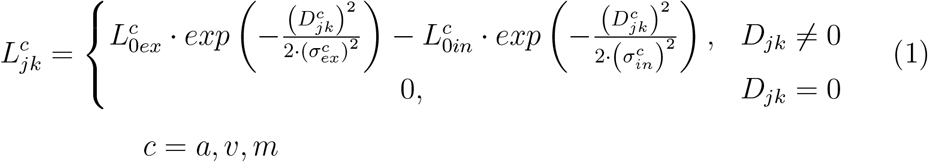

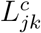 denotes the weight of the synapse from the pre-synaptic neuron at position *k* to post-synaptic neuron at position *j* within the area *c* (auditory, visual or multisensory). 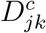 indicate the distance between the presynaptic neuron and the postsynaptic neurons. The excitatory Gaussian function is defined by the parameters 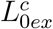 and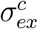, whereas the inhibitory function is defined by 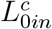 and 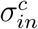. A null term (i.e. zero) was included in equation 1 to avoid auto-excitation.

#### 2.4.2. Cross-modal processing deficits

We modelled early cross-modal processing deficits by increasing or decreasing the weights of cross-modal connectivity. This was implemented by manipulating the value of the parameter 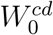 in both sensory modalities, defined in the equation 2:

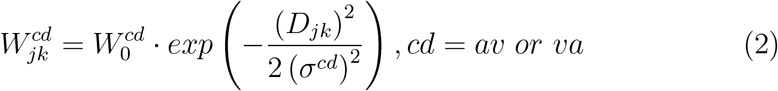

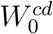 denotes the highest level of synaptic efficacy and *D*_*jk*_ is the distance between the neuron at position *j* in the postsynaptic unisensory region and the neuron at position *k* in the presynaptic unisensory region. *σ*^*cd*^ defines the width of the cross-modal synapses. These synaptic weights are symmetrically defined (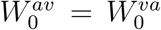 and *σ*^*av*^ = *σ*^*va*^) by the Gaussian function described above.

#### 2.4.3. Failures in top-down signalling

The failures in top-down signalling were implemented by uniformly weakening top-down synaptic weights. This was achieved by changing the value of the parameter 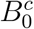, defined in equation 3:

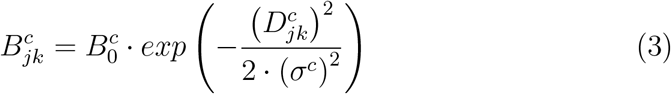

Here, 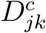 is the distance between the multisensory neuron at position j and the unisensory neuron at position k. 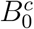 denote the value of the feedback when 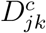 is equal to zero, representing the highest synaptic efficacy. *σ*^*c*^ represents the width of the feedback synapses.

#### 2.4.4. Decrease of synaptic density

Following Hoffman and Dobscha (1989), Hoffman and McGlashan (2006) and Paredes et al. (2022), the decrease in synaptic density between multisensory and unisensory neurons was implemented by resetting the connection weights of the feedforward synapses that were below a certain threshold 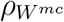 to zero. Similarly, the connection weights of the cross-modal synapses with weights below 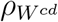 were reset to zero, to decrease the density of the crossmodal synapses between unisensory areas.

## 3. Results

### 3.1. Parameter exploration

We systematically varied the parameters governing the strength of lateral excitatory (*L*_0*ex*_), feedback 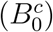 and cross-modal 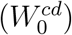 weights, and feedforward 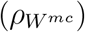 and cross-modal 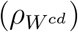 pruning to explore plausible mechanisms behind the changes in the temporal window of illusion (TWI) observed in individuals with H-SPQ (see Figure 3A). Our simulations revealed that an increase in recurrent excitation weights enlarges the temporal window of illusion. Changes in feedback weights or pruning feedforward synapses had a weaker effect.

**Figure 3:**
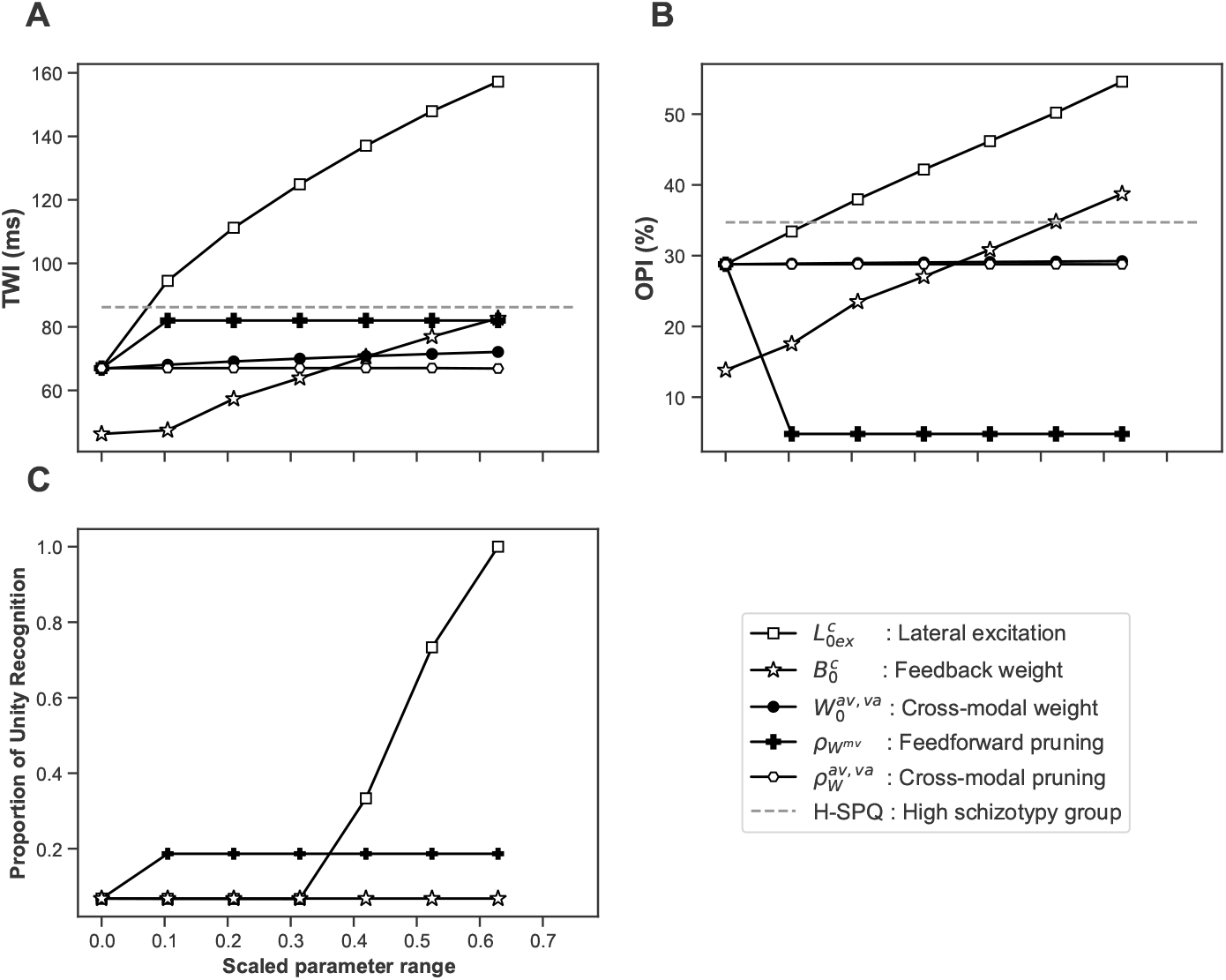
Effects of potential neural impairments in the temporal window of illusion, overall proneness to the illusion and causal inference in the network model. **Panels A, B** and **C** show the effect of systematic variation of parameters in the range at which they produce sensible responses to stimuli (these ranges were scaled to facilitate visual comparison). The dashed lines represent the average responses of H-SPQ individuals in the experimental study (Ferri et al., 2018). **Panel A** shows that an increase in the temporal window of illusion observed in H-SPQ could be reproduced by an increase in lateral excitation 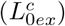. **Panel B** indicates that the overall proneness to experience the double flash illusion observed in H-SPQ could be reproduced by an increase in lateral excitatory 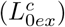 or feedback weights 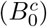. **Panel C** indicates that both an increase in lateral excitation 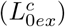 weights or pruning feedforward synapses 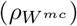 increases the overall probability of inferring a common cause to the presented stimuli.

Similarly, we found that an increase in lateral excitatory or feedback weights leads to an increase in the overall proneness to experience the illusion (OPI) (i.e. the percentage of reports of perceiving two flashes across all SOAs) at the level observed in individuals with H-SPQ (see Figure 3B). The opposite effect was observed when manipulating the pruning of feedforward synapses.

Furthermore, we varied these same parameters to explore their impact on the causal inference mechanism of the model. Our simulations revealed that an increase in lateral excitatory weights or pruning feedforward synapses increases the average probability of inferring a single common cause to the stimuli across all SOAs (see Figure 3C). Changes in cross-modal connectivity did not affect causal inference and are excluded from the figure.

### 3.2. Identification of the H-SPQ model

To select the model that provides the best fit to the data of the H-SPQ group (Ferri et al., 2018), we considered the Root Mean Square Error (RMSE) adjusted by the number of free parameters, as in Paredes et al. (2022).

The model that best fit the data from the H-SPQ group is presented in Figure 4A. Our findings indicate that increased lateral excitation in unisensory and multisensory areas matches H-SPQ responses in the DFI task (adj RMSE = 2.55). We discarded models with increased cross-modal (adj RMSE = 7.43) or feedback weights (adj RMSE = 5.89), or feedforward pruning (adj RMSE = 7.94). The model responses stay within participants’ standard error range, except for SOAs under 75 ms (see Supplementary Information). The TWI from our L-SPQ and H-SPQ models approximate the observed data, as shown in Figure 4B.

**Figure 4:**
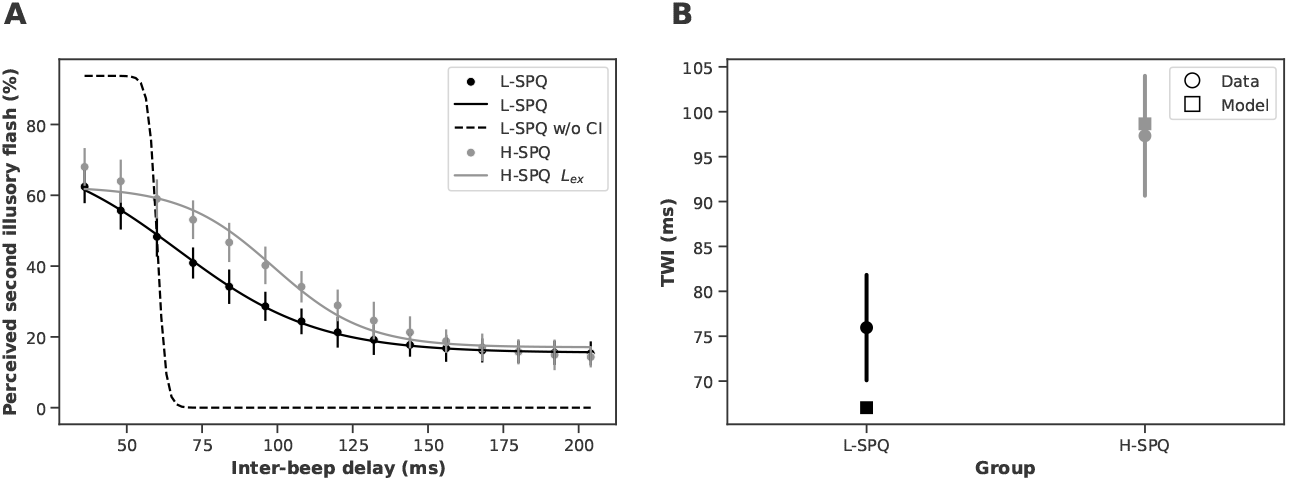
L-SPQ and H-SPQ network models identified by the fitting procedure. **Panel A** shows the DFI task responses generated by the L-SPQ and H-SPQ network models. The solid lines depict the sigmoid fit obtained from the data generated by the models. The dots represent the discretisation of the sigmoid fit obtained from the data collected in the experiment (Ferri et al., 2018). The quantitative fitting of the L-SPQ model (black) using 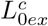 (grey) is sufficient to reproduce the H-SPQ data. The black dashed lines represent an alternative L-SPQ model without the causal inference area. **Panel B** shows the temporal window of illusion (TWI) generated by the L-SPQ and H-SPQ network models. Squares depict the TWI generated by the models, whereas dots and standard error bars depict the TWI observed in the experiment Ferri et al. (2018).

## 4. Discussion

This study aimed at modelling the neural mechanisms behind the higher proneness to the double flash illusion observed in individuals scoring high in schizotypy (Ferri et al., 2018). For this purpose, an existing double flash illusion audiovisual network model (Cuppini et al., 2014) was adapted to incorporate a multisensory layer with feedforward and feedback connectivity to unisensory layers, allowing iterative causal inference across the network (Rohe and Noppeney, 2015a; Rohe et al., 2019). This adaptation allowed us to quantitatively fit this model to behavioural data for the first time, revealing the crucial role of feedback input to generate sound-induced visual illusions (see Figure 2).

We examined how E/I imbalance, cross-modal processing deficits, topdown signalling failures, and synaptic decrease affect illusory responses. These neural impairments align with NMDA receptor and GABA neuron dysfunctions, along with heightened D_2_ receptor activity, crucial in schizophrenia models (Sterzer et al., 2018; Lanillos et al., 2020; Leptourgos et al., 2022). Analysing the experimental data (Section 3.2), we identified a neural mechanism related to a higher susceptibility to the double flash illusion in high schizotypy individuals: increased recurrent excitation in unisensory and multisensory neurons. Simulations (Section 3.1) show that this mechanism expands the illusion’s temporal window (TWI) and raise overall proneness (OPI), also increasing the probability that the network infers a common cause from stimuli.

### 4.1. An E/I imbalance account of reduced temporal discrimination

Our results are broadly in line with magnetic resonance spectroscopy (MRS) studies that consistently show alterations in glutamatergic and GABAergic neurotransmission in the schizophrenia spectrum (Reddy-Thootkur et al., 2022). Particularly, a disrupted E/I balance has been associated with a wider temporal window of illusion in individuals with schizotypal traits: concentrations of glutamatergic compounds (Glx) mediated by individual E/I genetic profiles predict multisensory temporal discrimination in the Simultaneity Judgement (SJ) task and cognitive-perceptual scores on the SPQ scale. Higher concentrations of Glx during the SJ task correlate with lower temporal discrimination and higher schizotypy in individuals with a genetic shift toward greater excitation (Ferri et al., 2017).

Nevertheless, our model reproduces the E/I balance perturbation but does not distinguish between specific impairments in excitation or inhibition (e.g. failures in GABAergic or glutamatergic receptors). The model could be expanded to account for differences in inhibition by adopting a modelling approach based on spiking neurons and the direct manipulation of NMDA and GABA input currents (e.g. Abbasi et al. (2023)). This would enable the possibility link our results with a recent animal study showing that a reduction in GABAergic inhibition in the audiovisual cortex is sufficient to alter multisensory processing and reduce audiovisual temporal discrimination (Schormans and Allman, 2023). Here, wild-type rats received Gabazine (a GABA-A receptor antagonist) in the V2L cortex (an area known to process audiovisual stimuli) and evaluated during audiovisual stimulation and the Temporal Order Judgement (TOJ) task. The infusion of gabazine was sufficient to widen the temporal window of audiovisual integration and decrease the perception of time differences between auditory and visual stimuli.

Future work could also extend the model to better describe neural dynamics and explore how changes in E/I balance influence oscillatory patterns in multisensory integration areas (Fotia et al., 2021). Electrophysiological evidence has consistently shown a reduction in the power of gamma band activity in patients with SCZ (Leicht et al., 2010; Lenz et al., 2011; Leicht et al., 2015; Keil et al., 2016). In the DFI paradigm, this is observed as a lack of enhancement of beta/gamma band (25-35 Hz) responses in trials where the illusory flash is perceived, suggesting early binding deficits in SCZ (Balz et al., 2016b). Such an approach could enable the computational understanding of how the concentration of GABA in STS is correlated with the perception of DFI and influences the oscillatory activity of the gamma band (Balz et al., 2016a).

To our knowledge, this is the first study to show computationally at the network level how a biologically plausible deregulation of the E/I balance reduces temporal sensitivity (indexed by the size of the TWI) in the schizophrenia spectrum. Moreover, we show how this impairment causes a higher proneness to experience sound-induced visual illusory phenomena (Ferri et al., 2018; Haß et al., 2017). In line with Paredes et al. (2022), we suggest that this mechanism causes the decreased sensitivity observed in schizophrenia and high schizotypal individuals in variants of the task (e.g., tactile-induced double flash illusion) (Fotia et al., 2021) and related spatiotemporal discrimination paradigms (Tseng et al., 2015; Ferri et al., 2016; Zhou et al., 2018; Dalal et al., 2021; Di Cosmo et al., 2021).

### 4.2. Possible links with Bayesian accounts of SCZ

Our modelling approach could be seen as a way to bridge the neural and Bayesian level of analysis of sound-induced visual illusory phenomena. Our results expand previous network modelling of audiovisual integration showing that cross-modal synapses encode prior probabilities of the co-occurrence of audiovisual stimuli in spatial localisation tasks (Ursino et al., 2017, 2019). This view aligns with Bayesian Causal Inference modelling of the Simultaneity Judgement task, indicating that temporal binding windows are influenced by an a priori bias to bind sensory information (Magnotti et al., 2013; Noel et al., 2018a; Chancel et al., 2022).

Future research could aim to link our results to Bayesian accounts of schizophrenia (Friston et al., 2016; Jardri et al., 2016; Sterzer et al., 2018; Tarasi et al., 2022a,b, 2023). A recent Bayesian model of the sound-induced flash illusion suggests that wider temporal binding windows and higher illusion rates would be a consequence of reduced precision in unisensory modalities (Zhu et al., 2024). However, it is unclear how our findings on increased excitation in recurrent connectivity within the multisensory integration network align with Bayesian accounts of psychotic symptoms: strong priors (Teufel et al., 2015; Powers et al., 2017; Cassidy et al., 2018); weaker or variable priors (Nazimek et al., 2012; Samaha and Romei, 2024; Schmack et al., 2015, 2017; Jardri et al., 2017; Valton et al., 2019; Fletcher and Teufel, 2022; Goodwin et al., 2023); stronger priors at higher sensory processing levels with weaker priors at lower levels (Yang et al., 2016; Petrovic and Sterzer, 2023); or circular inference in multisensory integration (Leptourgos et al., 2022). We encourage direct evaluation of the activity of the unisensory and multisensory areas during sound-induced flash illusion in patients with schizophrenia to identify whether aberrations in top-down or bottom-up processing drive the increased proneness to visual illusory phenomena.

### 4.3. Multisensory causal inference in bodily self-aberrations

To our knowledge, this is the first study to computationally examine explicit multisensory causal inference in the schizophrenia spectrum (French and DeAngelis, 2020). We showed at the network level how an increase in excitation in recurrent connectivity potentially increases the overall probability of inferring a common cause from sensory stimuli (see Figure 3C).

Speculatively, our predictions about multisensory causal inference impairments due to increased excitation could be related to bodily self-aberrations observed in the SCZ spectrum (Klaver and Dijkerman, 2016; Michael and Park, 2016; Sandsten et al., 2020; Di Cosmo et al., 2021). Empirical evidence consistently shows a co-occurrence of reduced spatiotemporal discrimination and impaired body ownership indexed by a higher proneness to experience the Rubber Hand Illusion (RHI) in the SCZ spectrum (Rossetti et al., 2020; Costantini et al., 2020; Zopf et al., 2021; He et al., 2022).

We tentatively suggest that an increase in excitatory weights of recurrent synapses within multisensory integration networks increases susceptibility to RHI and bodily perceptual symptoms observed in the SCZ spectrum (Laurin et al., 2021; He et al., 2022; Torregrossa and Park, 2022). We encourage further research into the neurobiology of causal inference in SCZ during the RHI and other multisensory illusions (Klaver and Dijkerman, 2016; Michael and Park, 2016; Ferroni et al., 2019; Sandsten et al., 2020) to better understand bodily self-disturbances in patients with SCZ.

### 4.4. Limitations and Future Directions

The network model used in this study is a rough simplification of electrochemical processes and synaptic connectivity of neurons that compute multisensory integration. As a consequence, our network does not account for the dynamics of oscillatory activity observed in illusory phenomena induced by sound or touch (Balz et al., 2016a,b; Fotia et al., 2021). Moreover, our network does not accurately reproduce the responses observed at lower inter-beep delays (see Supplementary Information). At best, the network model presented here serves as a tool for theoretical reasoning that can be incrementally improved for biological realism (Guest and Martin, 2023).

Furthermore, our modelling approach does not consider relevant neural input from frontal areas. This input is crucial considering that cognitive load and attention modulate the proneness to experience the DFI (Keil, 2020) and specific impairments in selective attention and cognitive control have been observed in the schizophrenia spectrum (Gold et al., 2007; Lesh et al., 2011). Furthermore, neuroimaging evidence suggests a critical role for frontotemporal regions in multisensory integration impairments observed in SCZ (Gröhn et al., 2022; Leptourgos et al., 2022).

The experimental evidence directly evaluating sound-induced flash illusions in the SCZ spectrum has only begun to accumulate (Vanes et al., 2016; Haß et al., 2017; Balz et al., 2016b; Ferri et al., 2018). This study focuses on temporal binding without ruling out other factors contributing to temporal distortions in schizophrenia, like context integration gain (Cohen et al., 1999) or prediction delay mechanisms (Whitford et al., 2012; Okimura et al., 2023). Our findings are based on modelling a study with healthy individuals without evaluating causal inference (Ferri et al., 2018). Various experimental paradigms explore sound-induced flash illusions with different stimuli and spatiotemporal manipulations (Hirst et al., 2020; Keil, 2020). These paradigms remain underexplored in the SCZ spectrum, complicating the understanding of illusion proneness in SCZ (Vanes et al., 2016). Further studies with diverse stimuli and clear causal inference and spatiotemporal sensitivity measurements in the SCZ spectrum are needed (Klaver and Dijkerman, 2016; Michael and Park, 2016; Ferroni et al., 2019; Sandsten et al., 2020).

## 5. Funding

F.F. is supported by the “Departments of Excellence 2023–2027” initiative of the Italian Ministry of Education, University and Research for the Department of Neuroscience, Imaging and Clinical Sciences (DNISC) of the University of Chieti-Pescara, and by the Italian Ministry of University and Research (MUR), funded by the European Union – NextGenerationEU, under the National Recovery and Resilience Plan (NRRP) CUP: D53D23020890001. V.R. is supported by MUR – Ministry of University and Research, Italy (P2022XAKXL and 2022H4ZRSN), Ministerio de Ciencia, Innovación y Universidades, Spain (PID2019111335GA-100) and BIAL Foundation (033/22).

## 6. CRediT authorship contribution statement

**Renato Paredes:** Conceptualisation, Methodology, Software, Formal Analysis, Writing Original Draft. **Francesca Ferri:** Conceptualisation, Investigation, Writing - Review and Editing. **Vincenzo Romei:** Conceptualisation, Investigation, Writing - Review and Editing. **Peggy Seriés:** Conceptualisation, Methodology, Supervision, Writing - Reviewing and Editing.

## 7. Declaration of competing interest

None.

## 8. Availability of data and materials

The model was implemented using the Scikit-NeuroMSI framework for modelling multisensory integration (Paredes et al., 2023). The code used to produce the simulations presented in this manuscript can be found at: https://github.com/renatoparedes/SCZDFImodel.

## 9. Acknowledgements

We acknowledge Denise Robles Soberón (Hey Cometa) for her contribution in the graphics employed in this paper.

## Supplementary Information

### 1 Participants

We modelled the data collected in the study conducted by Ferri et al. (2018). One hundred and ninety-six adult volunteers were screened for schizotypal traits using the Schizotypal Personality Questionnaire (SPQ) (Raine, 1991). The questionnaire was administered online through Qualtrics. Participants were categorized into quintiles based on their SPQ scores. 64 participants participants falling within the first and fifth quintile were selected: 32 with low schizotypy scores (range: 5-16) and 32 with high schizotypy scores (range: 36-68). All participants had normal or corrected vision and hearing and no history of substance abuse or other psychiatric disorders. Age and gender were matched between the two groups. Demographic details are presented in Table 1.

**Table 1:** Demographics.

| Trait | L-SPQ | H-SPQ |
| --- | --- | --- |
| SPQ Score Range | 5-16 | 36-68 |
| No. of Females | 25 | 25 |
| Mean Age (SD) | 29 (9) | 26 (7) |
| British White | 41% | 44% |
| Other White | 50% | 50% |
| Asian/Indian | 9% | 6% |

Three participants in the low schizotypy group and four in the high schizotypy group were excluded from the analysis due to unreliable responses. This resulted in a final sample of 57 participants: 29 in the low schizotypy group and 28 in the high schizotypy group. All participants provided written informed consent before participating in the study.

### 2 Temporal Window of Illusion

Following Ferri et al. (2018), to determine the Temporal Window of Illusion (TWI), we analysed the proportion of trials in which participants reported perceiving the DFI (i.e., “two” responses) at each Stimulus Onset Asynchrony (SOA). For each participant, we plotted the proportion of DFI reports against the SOA and fitted a psychometric sigmoid function to these data points:

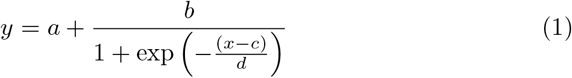

where:

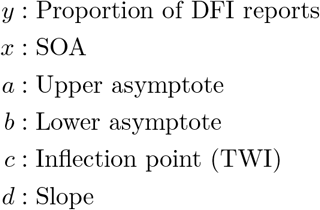

The estimated SOA (ms) corresponding to the inflection point (c) of the sigmoid function was defined as the individual’s TWI, representing the temporal window within which the illusion was maximally experienced.

### 3 Multisensory causal inference network model

The network model used in this study is an extension of previous models developed by Cuppini and his collaborators (Cuppini et al., 2014, 2017). The model consists of three layers: two encode auditory and visual stimuli, separately, and connect to a multisensory layer where causal inference is computed. Each of these layers consists of 30 neurons arranged topologically to encode a 30° space. Hence, each neuron encodes 1° of space and neurons close to each other encode close spatial positions.

Each neuron will be noted with a superscript *c* indicating a specific cortical area (a, v or m for the auditory, visual or multisensory area respectively). Similarly, each neuron will hold a subscript *j* refereed to its spatial position within a given area.

Neurons in each layer have a sigmoid activation function and first-order dynamics:

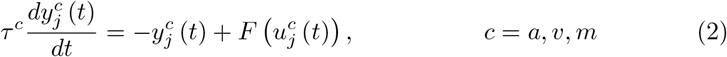

Here, *u*(*t*) and *y*(*t*) are used to represent the net input and output of a given neuron at time *t. τ*^*c*^ denotes the time constant of neurons belonging to a given area *c. F* (*u*) represents the sigmoidal relationship:

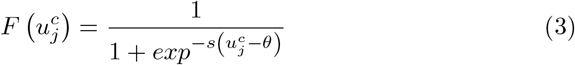

Here, *s* and *θ* denote the slope and the central position of the sigmoidal relationship respectively. Neurons in all regions differ only in their time constants, chosen to mimic faster sensory processing for stimuli in the auditory region compared to visual stimuli.

The net input of a neuron is the sum of an inside (i.e. within region) component 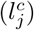 and an outside (i.e. extra-area) component 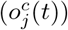:

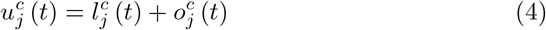

The within region component 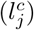 is defined as:

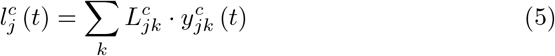

Here 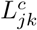 represents the strength of the lateral synapse from a presynaptic neuron at position *k* to a postsynaptic neuron at position *j* in the region *c*. 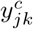 is the activity of the presynaptic neuron at position *k*.

Such synapses are symmetrical and arranged according to a “Mexican hat” pattern (i.e. a central excitatory area surrounded by an inhibitory ring for each neuron, so that the entire layer generates excitation for spatially close stimuli and inhibition for distant stimuli):

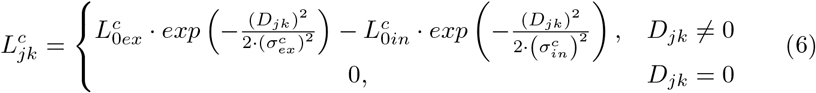

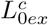 and 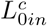 denote the highest level of excitatory and inhibitory synaptic efficacy in the region *c*, respectively. *D*_*jk*_ indicate the distance between the pre-synaptic neuron and the post-synaptic neurons within a given area:

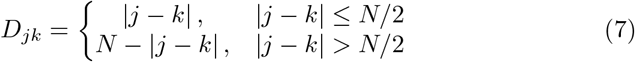

This defines a circular structure where each neuron receives the same number of lateral connections.

Importantly, the extra-area input is defined differently for unisensory and multisensory areas. The extra-area input for the unisensory areas includes a stimulus from the external world 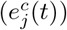, a cross-modal component coming from the other unisensory area 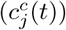, a feedback component coming from the multisensory area 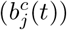 and a noise component 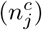.

The stimulus from the external world is simulated as a 1-D Gaussian function to represent the uncertainty in the detection of external stimuli:

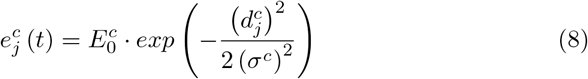

Here, 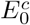 denotes the strength of the stimulus, 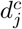 the distance between neuron at position *j* and the stimulus at position *p*^*c*^, and *σ*^*c*^ the degree of uncertainty in sensory detection. The distance 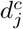 is defined as:

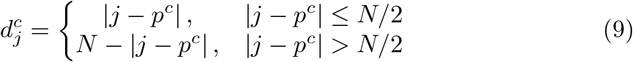

Furthermore, the cross-modal input is defined as:

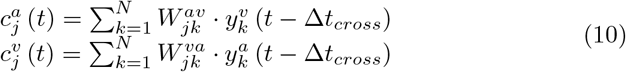

Here Δ*t*_*cross*_ represents the latency of cross-modal inputs between two unisensory regions. The synaptic weights are symmetrically defined (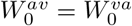 and *σ*^*av*^ = *σ*^*va*^) by the Gaussian function:

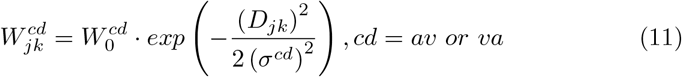

*W*_0_ denotes the highest level of synaptic efficacy and *D*_*jk*_ is the distance between neuron at position *j* in the post-synaptic unisensory region and the neuron at position *k* in the pre-synaptic unisensory region. *σ*^*cd*^ defines the width of the cross-modal synapses.

Furthermore, the feedback input is defined as:

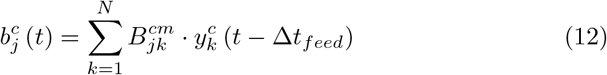

Here Δ*t*_*feed*_ represents the latency of feedback inputs between the multisensory and unisensory regions. The feedback synaptic weights are also symmetrically (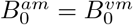 and *σ*^*am*^ = *σ*^*vm*^) defined:

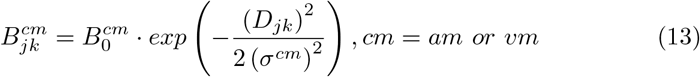

Here, 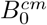 denotes the highest level of synaptic efficacy and *D*_*jk*_ is the distance between neuron at position *j* in the post-synaptic unisensory region and the neuron at position *k* in the pre-synaptic multisensory region. *σ*^*cd*^ defines the width of the feedback synapses.

The noise component 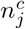 is extracted from a standard uniform distribution on the interval [*n*_*max*_ + *n*_*max*_]. Here *n*_*max*_ is defined as the 40% of the strength of the external stimulus for each modality.

All these external sources are filtered by a second order differential equation to mimic the temporal dynamics of the stimuli in a cortex:

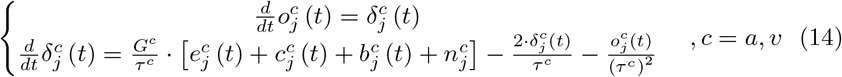

Here, *G*^*c*^ represents gain and *τ*^*c*^ the time constants of the dynamics.

In contrast, the extra-area input for the multisensory area comes only from feedforward synapses from the two unisensory areas:

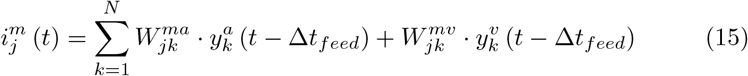

Here Δ*t*_*feed*_ represents the latency of feedforward inputs between the unisensory and multisensory regions. 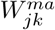 and 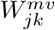 are the synapses connecting the pre-synaptic neuron at position *k* in a given unisensory area and the postsynaptic neuron at position *j* in the multisensory area.

The weights of these feedforward synapses are symmetrically defined as:

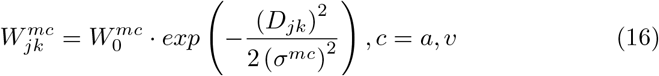

Here 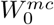 denotes the highest value of synaptic efficacy, *D*_*jk*_ the distance between the multisensory neuron at position *j* and the unisensory neuron at position *k*, and *σ*^*mc*^ the width of the feedforward synapses.

These external sources are also filtered by a second order differential equation:

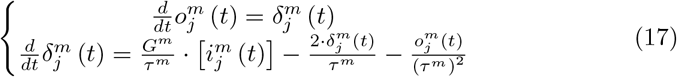

Here, *G*^*m*^ represents gain and *τ*^*m*^ the time constants of the dynamics in the multisensory neurons.

### 4 Values of L-SPQ model parameters

### 5 Parameter exploration

We systematically varied the parameters governing recurrent excitation (*L*_0*ex*_), the strength of feedback 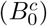 and cross-modal 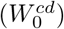 weights, and feedforward 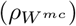 and cross-modal 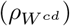 pruning to explore plausible mechanisms behind the changes in the temporal window of illusion (TWI) observed in individuals with H-SPQ (see Figure 3).

In this parameter exploration, we examined seven uniformly distributed values within the operational range of each aforementioned parameter. The operational range is characterized by the parameter variations for which the base model (L-SPQ) demonstrates responses that align with those observed in human participants during the behavioral experiment (Ferri et al., 2018). The bounds considered for the parameter exploration for 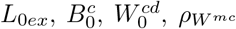, and 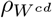 were (0.5, 0.65), (0.24, 0.25), (0.01, 0.055), (0, 0.35), and (0, 0.013), respectively.

### 6 Fitting procedure

The probability of perceiving two flashes in a single trial *P* (2*f*) was defined as the product of the two peak values above .15 read from visual neurons. We used a fitting procedure with the cost function defined by Equation 18.

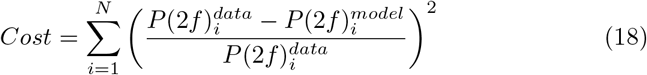

Here, 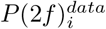 and 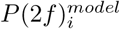 denote the P (2f) measured in the *i* th stimulus onset asynchrony (SOA). N represents the number of SOAs measured (i.e. 15 in the empirical study).

This cost function was minimised by the implementation of the differential evolution algorithm (Storn and Price, 1997) available in the SciPy v1.11.4 library for the Python programming language (Virtanen et al., 2020).

*P* (2*f*)^*model*^ is composed of the 15 *P* (2*f*) generated by the model at each SOA defined in the experimental task. Similarly, *P* (2*f*)^*data*^ is composed of 15 proportion values taken from the mean sigmoid fit obtained out of the experimental data of a given group. We assumed that the percentage of perceived second illusory flash reported in the original study Ferri et al. (2018) is equivalent to the probability of perceiving two flashes in each condition.

#### 6.1 L-SPQ Fitting

We fitted the model to the L-SPQ group data. Here, 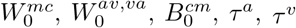 and *τ*^*m*^ (see Equations 16, 11, 13 14 and 17) varied freely to represent the group proneness to the illusion due to stable anatomical factors (i.e. immutable throughout the duration of the experiment). All other parameters (see Table 2) remained fixed and set to the values published by Cuppini et al. (2014, 2017). The bounds given to the optimisation algorithm to find 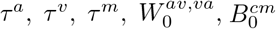 and 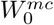, were (6, 20), (6.25, 60), (6, 120), (0.001, .05), (0.001, 1) and (0.001, 15) respectively. The results of the fit to the data of the L-SPQ group are presented in Table 2.

**Table 2:** Values of the parameters in the L-SPQ model. This parameterisation remained fixed throughout the simulations computed to identify the H-SPQ model.

|  |  |  |  |
| --- | --- | --- | --- |
| Neurons |  |  |  |
| $N = 30$ | $\theta = 20$ | $s = 0.3$ | $\tau = 1$ |
| External stimuli |  |  |  |
| $E_0^a = 2.325$ | $\sigma^a = 32^\circ$ | $E_0^v = 1.45$ | $\sigma^v = 4^\circ$ |
| Input dynamics |  |  |  |
| $G^{a,v,m} = e^1$ | $\tau^a = 6.56$ | $\tau^v = 9.19$ | $\tau^m = 120$ |
| $\Delta t_{cross} = 16 \text{ ms}$ | $\Delta t_{feed} = 95 \text{ ms}$ | | |
| Lateral synapses |  |  |  |
| $L_{0ex}^c = 0.5$ | $L_{0in}^c = 0.4$ | $\sigma_{ex}^c = 3^\circ$ | $\sigma_{in}^c = 24^\circ$ |
| Inter-areal synapses |  |  |  |
| $W_0^{mc} = 3.892$ | $\sigma^{mc} = 0.5^\circ$ | $B_0^{cm} = 0.623$ | $\sigma^{cm} = 0.5^\circ$ |
| $W_0^{av,va} = 0.001$ | $\sigma^{av,va} = 5^\circ$ | | |

We also fitted an alternative L-SPQ model without the causal inference area. We maintained the same base parameterisation, and used the optimisation algorithm to find *τ*^*a*^, *τ*^*v*^ and 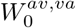. The bounds given were (6, 20), (6.25, 60), (6, 120) and (0.001, .05). This latter model failed to reproduce responses to SOAs beyond 60 ms, as shown in Figure 4A.

#### 6.2 H-SPQ Fitting

We fitted the L-SPQ model to the H-SPQ data. The bounds given to the optimisation algorithm to find 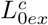 and 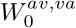 were (1.5, 1.7) and (0, 0.06) respectively. These limits were chosen because the response profile generated by values outside of this range ceases to resemble a sigmoid curve, which is characteristic of the average response profile found in the empirical study (Ferri et al., 2018).

It is observed that the model may not accurately reproduce the DFI responses observed at very short inter-beep intervals (below 75 ms). This discrepancy may be attributed, at least in part, to the fixed assumptions pertaining to the temporal delays of cross-modal and feedforward-feedback inputs incorporated within the model. Additionally, it is essential to acknowledge that the literature indicates substantial variability in DFI responses at very short inter-beep intervals across different individuals (Hirst et al., 2020). We recognize that further refinement of the model’s parameters and architecture may be requisite to more precisely capture the complexities of DFI perception over the entire spectrum of inter-beep intervals.

## Footnotes

1 Explicit and implicit causal inference refers to tasks in which participants have to directly evaluate the causal relationship of the stimuli (unity judgement) or indirectly by providing estimates of their spatiotemporal location (Acerbi et al., 2018).

2 The combination of unisensory and multisensory estimates weighted by the posterior probabilities of common and independent cause models.

3 When more than two visual peaks exceed the threshold, the average of all possible product combinations of these peaks is calculated.

## Notes

https://github.com/renatoparedes/SCZ_DFI_Model

### Competing Interest Statement

The authors have declared no competing interest.

### Summary of Updates

New simulations were performed to confirm the results and compare alternative models. This new version includes changes mainly in the results and discussion sections.

